# Self-assembly and contraction of micron-scale DNA rings

**DOI:** 10.1101/2023.03.09.531887

**Authors:** Maja Illig, Kevin Jahnke, Marlene Scheffold, Ulrike Mersdorf, Hauke Drechsler, Stefan Diez, Kerstin Göpfrich

## Abstract

Contractile rings formed from cytoskeletal filaments mediate the division of cells. The reverse-engineering of synthetic contractile rings could shed light on fundamental physical principles of the ring self-assembly and dynamics independent of the natural protein-based compounds. Here, we engineer DNA nanotubes and crosslink them with a synthetic peptide-functionalized star-PEG construct. The star-PEG construct induces the formation of DNA nanotube bundles composed of several tens of individual DNA nanotubes. Importantly, the DNA nanotube bundles curve into closed micron-scale DNA rings in a high-yield one-pot self-assembly process resulting in several thousand rings per microliter. The crosslinked DNA rings can undergo contraction to less than half of their initial diameter by two distinct mechanisms, triggered by increasing molecular crowding or temperature. DNA-based contractile rings expand the toolbox of DNA nanotechnology and could be a future element of an artificial division machinery in synthetic cells.

## Introduction

Cell division is a hallmark of life. After duplication and separation of the genetic information, the cellular compartment has to be divided to give rise to two daughter cells. Nature’s solution for compartment division is the formation of contractile rings made from cytoskeletal filaments (*1*). In eukaryotic cells, micronscale actomyosin rings assemble at the cell’s equatorial plane at the end of mitosis and meiosis (*2*). The ring contraction can be powered by two distinct mechanisms, namely molecular motor activity and diffusable actin crosslinkers. Whereas the first is an active, energy-consuming process, the latter mechanism mediates passive entropy driven contraction. Recently, it has been shown that passive filament crosslinkers can generate filament sliding and contractile forces, that are sufficient to antagonize motor protein action in microtubule (*3*) and actin networks (*4*).

Bottom-up synthetic biology has pursued the long-term goal to reconstitute a minimal division machinery inside lipid vesicles to establish fundamental physical principles in isolation of the complex environment of a cell and to eventually engineer a self-replicating cellular system from scratch. Towards this goal, actomyosin rings have been formed in vitro (*5*) and their contraction has been demonstrated in confinement (*6*). In lipid vesicles, this led to membrane deformations (*7*). Nevertheless, the reconstitution of a division machinery which can complete the division of lipid vesicles remains an open challenge in bottom-up synthetic biology (*8, 9*). To tackle this challenge, it is important to critically ask which physical features are required for a minimal division system and how they contribute to the contraction process. Towards this end, it would be ground breaking to establish an entirely synthetic division machinery, which does not rely on nature’s building blocks. A fully engineered contractile ring could yield mechanistic insights into the biophysics of the process towards an alternative set of molecular components for the division of synthetic cells. There are ongoing efforts in the field of DNA nanotechnology to recreate functional mimics of cytoskeletal nanotubes from DNA. Of particular interest are DNA nanotubes (*10*), which are composed of individual DNA tiles assembled from five single-stranded DNA oligomers. Each DNA tile contains single-stranded DNA overhangs that allow for its polymerization into hollow, cylindrical nanotubes made of 12 DNA duplexes. These DNA nanotubes have been equipped with different features that mimic functions of a cytoskeleton, such as reversible assembly (*11, 12*), directional growth (*13*) and transport with engineered molecular motors (*14*) or enzyme activity (*15*). To mimic cellular confinement, DNA nanotubes have also been reconstituted in water-in-oil droplets (*15, 16*) and giant unilamellar lipid vesicles (GUVs) (*12*). Recently, we observed the formation of incomplete rings in GUVs upon addition of molecular crowders (*12*). However, no closed rings and no contraction could be obtained. Closed DNA rings have been assembled on the nanoscale from single-strands (*17*) or with DNA origami to template liposomes (*18*) or gold nanoparticles (*19*), to engineer liposome fusion and lipid transfer (*20*) or mechanically interlocked molecules (*21*) and they have been used as large-diameter membrane pores (*22*). However, these nanoscale DNA rings are one to two orders of magnitude too small to span the GUV circumference and it remains an open question how their contraction could be achieved.

Therefore, our aim is the self-assembly of DNA-based contractile rings on the micron scale. We reason that the assembly and contraction of micron-scale DNA nanotube rings can take inspiration from the mechanisms at play for nature’s cytoskeletons. While ring assembly clearly requires crosslinkers, contraction could either be achieved with suitable molecular motors or passive crosslinkers, in analogy to the contraction of protein-based cytoskeletal rings. Assuming that the passive crosslinker approach is more straight forward to adapt to DNA nanotubes, we require a synthetic crosslinker for DNA nanotubes. Multivalent positively charged peptides have successfully been shown to crosslink microtubules (*23*). Since both microtubules and DNA nanotubes have a negative surface charge, we hypothesize that similar crosslinkers also work for DNA nanotubes. Indeed, we show that we can bundle DNA nanotubes and achieve the self-assembly of closed micron-scale DNA rings in a one-pot bulk solution upon addition of multivalent positively charged peptides. We control the DNA bundle thickness as well as the ring diameter. Additionally, the contraction of the DNA rings to less than 45 percent of their initial diameter was achieved with these passive crosslinkers rather than active molecular motors. In the future, reverse-engineering the cell’s molecular hardware with DNA nanotechnology may be an alternative route to realize synthetic cell division.

## Results

### Synthetic peptides as crosslinkers for DNA nanotubes

We first assemble DNA nanotubes from the well-established double-crossover DNA tile design, whereby each tile consists of five DNA oligomers (Fig. 1**A**, Supporting Table S1) (*10*). Due to their sticky-end overhangs and their intrinsic curvature, these tiles assemble into hollow nanotubes consisting of 12 DNA duplexes. To form bundles and contractile rings from these DNA nanotubes, we need a synthetic crosslinker that satisfies the following requirements imposed by the nature of the DNA cytoskeleton: First, a crosslinker which binds to DNA nanotubes by electrostatic interactions has to be positively charged because of the negatively charged backbone of the DNA. Second, it has to be multivalent, so that it can crosslink multiple DNA nanotubes. Thus, we can make use of a multivalent positively charged peptide construct, which has already been shown to electrostatically interact with the negatively charged surface of microtubules thereby crosslinking them (*23*). The construct consists of a four-arm 10 kDa starPEG backbone which is coupled to seven lysine-alanine amino acid repeats on each of the four arms (Fig. 1**B**). We will refer to it as starPEG-(KA7)_4_. In a fully extended conformation, each arm would have a numerical length of approximately 30 nm. Hence, the construct exhibits four flexible arms with positively charged amino acid repeats that can bind to and crosslink the negatively charged DNA nanotubes by electrostatic interactions. To provide appropriate negative controls, we additionally synthesize monovalent KA7 peptides, which are not expected to cross-link DNA nanotubes, as well as a construct that features seven negatively charged aspartate-alanine repeats on a tetravalent starPEG backbone (starPEG-(DA7)_4_) which is not expected to bind to the negatively charged DNA backbone. All constructs are further labeled with 5-TAMRA (*λ*_*ex*_ = 561 nm) to allow for their visualization with confocal microscopy.

**Fig. 1.**
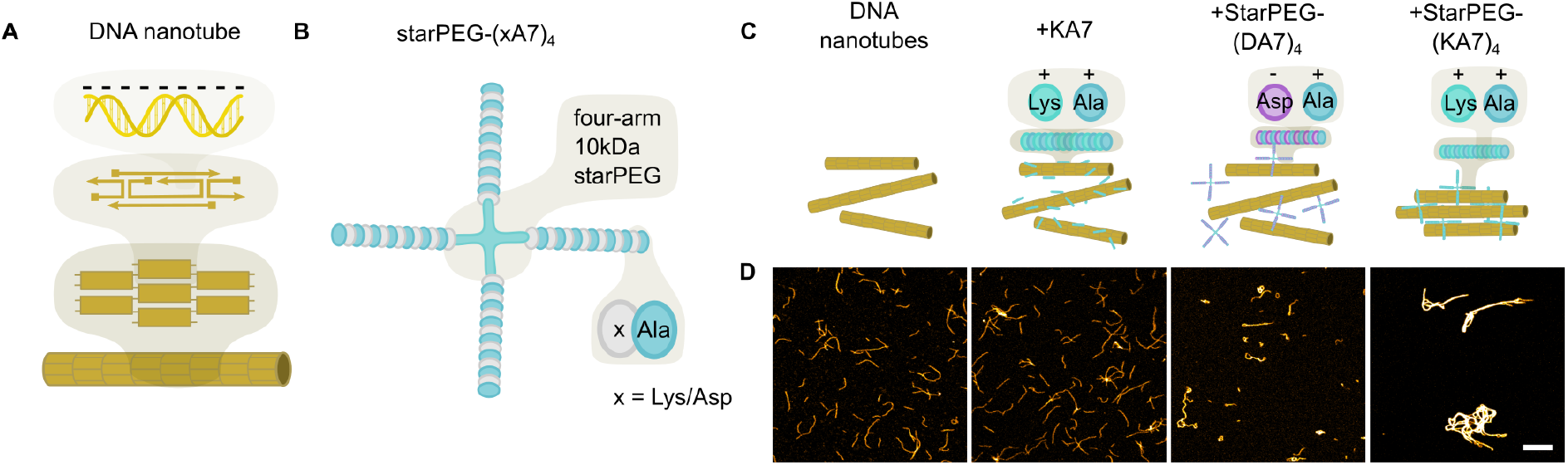
StarPEG-(KA7)_4_ bundles DNA nanotubes. (**A**) Schematic illustration of DNA nanotubes formed from double-crossover DNA tiles (*10*). (**B**) Schematic illustration of tetravalent starPEG-(xA7)_4_ composed of four branches of 7 lysine- or aspartate-alanine repeats. (**C**) Schematic illustration of DNA nanotubes in the absence and presence of different synthetic peptide construct. (**D**) Confocal images of DNA nanotubes (30 nM DNA tiles, labeled with Atto633, *λ*_*ex*_ = 640 nm) without any peptide; with 2 µM positively charged monovalent KA7-peptide; with 500 nM negatively charged tetravalent starPEG-(DA7)_4_ and with 500 nM positively charged tetravalent starPEG-(KA7)_4_ composed of four branches of 7 lysine-alanine repeats (from left to right). Scale bar: 10 µm.

To test whether starPEG-(KA7)_4_ indeed can bundle DNA nanotubes, we mix fluorescently-labeled DNA nanotubes with starPEG-(KA7)_4_ or the respective control constructs and image them by confocal microscopy (Fig. 1**C**,**D**). In presence of monovalent KA7 peptides, single DNA nanotubes remain homogeneously distributed across the observation chamber similar to the pure DNA nanotubes in absence of any peptide. Weak nanotube bundling can be observed in presence of the negatively charged starPEG-(DA7)_4_ (Fig. 1**D**), likely due to the positively charged magnesium-ions in solution. Notably, in presence of starPEG-(KA7)_4_, the DNA nanotubes form nanotube bundles with much higher fluorescence intensity extending to a length of several tens of micrometer (Fig. 1**D**, right image). We find that the extend of bundling correlates with the concentration of starPEG relative to the concentration of DNA tiles (Supplementary Fig. S1).

This shows how peptide engineering and DNA nanotechnology can be combined to form hybrid materials.

### Characterization of the DNA nanotube crosslinking with synthetic multivalent peptides

In particular, we need to establish whether the DNA nanotube bundling is caused by the depletion effect only, whereby starPEG-(KA7)_4_ acts as a molecular crowder, or whether the peptide can actually crosslink DNA nanotubes. We therefore immobilize DNA nanotubes on the surface of the observation chamber and preload them with starPEG-(KA7)_4_. We then wash out excess starPEG-(KA7)_4_ from the solution when adding a second type of DNA nanotubes, which we labelled with a different fluorophore in order to distinguish them from the immobilized DNA nanotubes (immobilized DNA nanotubes A: green; free DNA nanotubes B: yellow, Fig. 2**A**). After another washing step, only starPEG-(KA7)_4_ that is bound to the DNA nanotube A is present and responsible for the colocalization of the two DNA nanotube types (Fig. 2**B**).

**Fig. 2.**
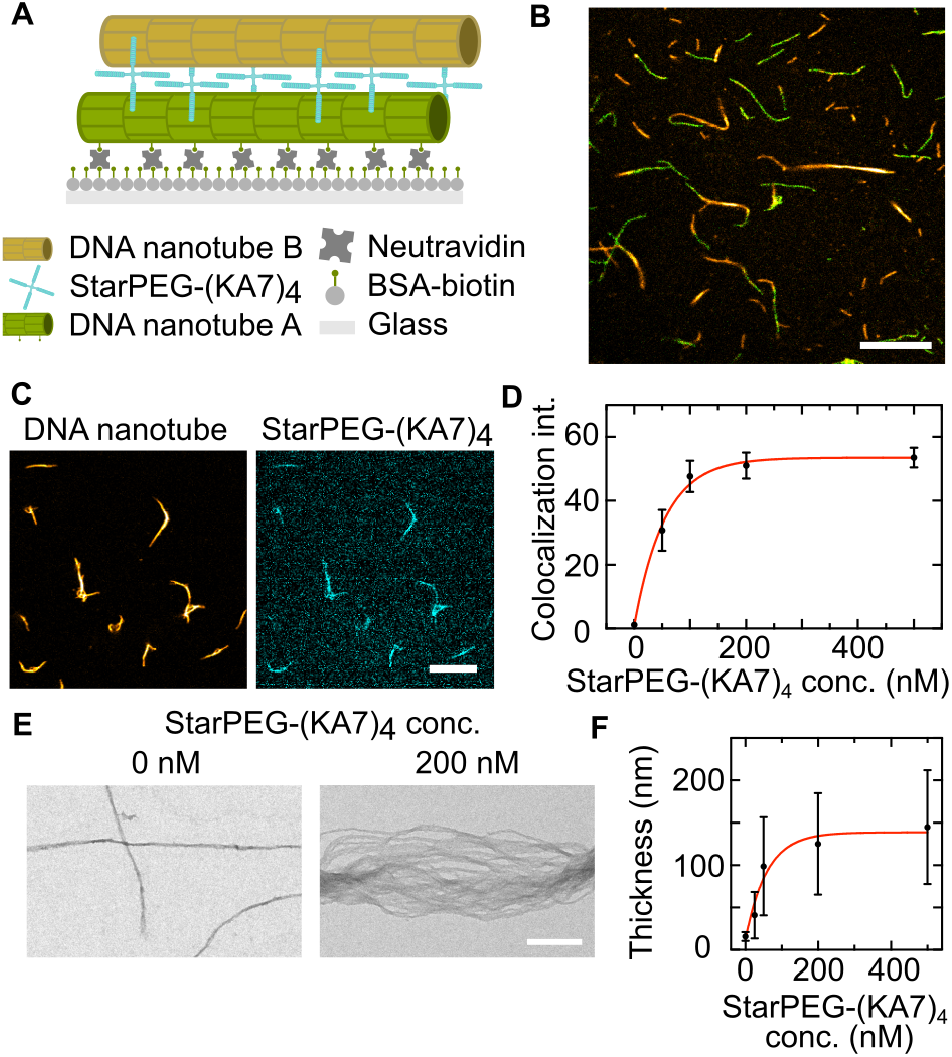
DNA nanotube colocalization and bundling by starPEG-(KA7)_4_. (**A**) Schematic illustration of the DNA nanotube colocalization assay with a biotinylated DNA nanotube A (green) bound to biotinylated bovine serum albumin-coated glass slides upon addition of ± 0.2 wt% neutravidin. StarPEG-(KA7)_4_ crosslinks DNA nanotube B (yellow) to the immobilized nanotube A. (**B**) Composite confocal image of the immobilized DNA nanotube A (green, 6-FAM, *λ*_*ex*_ = 488 nm) and DNA nanotube B (yellow, Atto633-labeled, *λ*_*ex*_ = 640 nm). Scale bar: 10 µm. (**C**) Confocal images of DNA nanotubes (30 nM DNA tiles, yellow, Atto633-labeled, *λ*_*ex*_ = 640 nm) and 200 nM starPEG-(KA7)_4_ (cyan, TAMRA-labeled, *λ*_*ex*_ = 561 nm). Scale bar: 10 µm. (**D**) Colocalization intensity of starPEG-(KA7)_4_ (TAMRA-labeled, *λ*_*ex*_ = 561 nm) with DNA nanotubes (30 nM DNA tiles) (Atto633-labeled, *λ*_*ex*_ = 640 nm, Mean ± SD, *n* = 10, exponential fit plotted as red line). (**E**) TEM micrographs of DNA nanotubes (30 nM DNA tiles) in the absence of starPEG-(KA7)_4_ (left) and in presence of 200 nM starPEG-(KA7)_4_ (right). Scale bar: 200 nm. (**F**) DNA nanotube bundle thickness over starPEG-(KA7)_4_ concentration (Mean ± SD, *n* = 100, 30 nM DNA tiles, exponential fit plotted as red line).

Thereby we conclude that the binding must be induced by crosslinking and not by the depletion effect only, as not enough starPEG-(KA7)_4_ is left in solution to cause crowding (Supplementary Fig. S1). To assess how much starPEG-(KA7)_4_ can bind to DNA nanotubes, we label the synthetic star-PEG peptides with fluorescent 5-TAMRA and analyze its colocalization with the DNA nanotubes (Fig. 2**C**), i.e. the fluorescence of starPEG-(KA7)_4_ colocalizing with the DNA nanotubes (see Experimental Section and Supplementary Fig. S2). The colocalization intensity saturates at around 200 nM starPEG-(KA7)_4_ for DNA nanotubes assembled from 30 nM DNA tiles (Fig. 2**D**, Supplementary Fig. S3). The concentration at which binding saturates agrees well with previous results for the microtubule binding of starPEG-(KA7)_4_ (*23*). Additionally, we analyze the DNA bundle thickness with negative stain transmission electron microscopy (TEM) for varying starPEG-(KA7)_4_ concentrations. Electron micrographs reveal the transition from single DNA nanotubes to DNA bundles consisting of tens of DNA nanotubes in presence of starPEG-(KA7)_4_ (Fig. 2**E**, Supplementary Fig. S4). Concomitantly, the bundle thickness increases by one order of magnitude from single DNA nanotubes with an apparent cross section of (15 ± 5)nm to bundles with a cross section of (145 ± 67)nm for 0 and 500 nM starPEG- (KA7)_4_, respectively. In accordance with the colocalization intensity, the bundle thickness also does not increase further than at 200 nM starPEG-(KA7)_4_ indicating the system’s maximal occupancy of starPEG-(KA7)_4_ on the DNA nanotubes (Fig. 2**F**). Dynamics in the time scale of the respecting experiments can be omitted since starPEG-(KA7)_4_ attaches to DNA nanotubes reaching saturation within minutes (Supplementary Fig. S5).

### Self-assembly of micron-scale DNA rings

To study the ability of starPEG-(KA7)_4_ to promote the self-assembly of DNA nanotubes into higher-order structures, we observe DNA nanotubes without immobilization in an unconstrained 3D environment, in solution. We find that starPEG-(KA7)_4_ promotes the efficient formation of DNA nanotube rings with several micrometres in diameter. Fig. 3**A** provides a confocal overview image that gives an impression of the high abundance of DNA rings, whereby the sample has been taken directly from the storage solution without additional concentration or purification steps. We obtain 5300 ± 2500 DNA rings per µL (Mean ± SD, Supplementary Fig. S6). StarPEG-(KA7)_4_ thus mimics the behaviour of septins in actin filament networks (*24*). The formation of closed rings is verified by TEM (Supplementary Fig. S7), which also reveals that the DNA rings typically consist of 10-50 nanotubes. We analyze the ring diameter for a range of starPEG-(KA7)_4_ to DNA tile ratios. For an excess of DNA nanotubes, the amount and the diameter of rings is significantly reduced compared to equimolar ratios (Fig. 3**B**,**C**). The ring diameter is, however, independent of the magnesium-ion concentration and the absolute concentrations of starPEG-(KA7)_4_ or DNA nanotubes (Supplementary Fig. S8). Finally, the abundance of DNA rings in a given sample volume is largely independent of time consistent with the fast attachment of starPEG-(KA7)_4_ to the DNA nanotubes (Supplementary Figs. S5, S6).

**Fig. 3.**
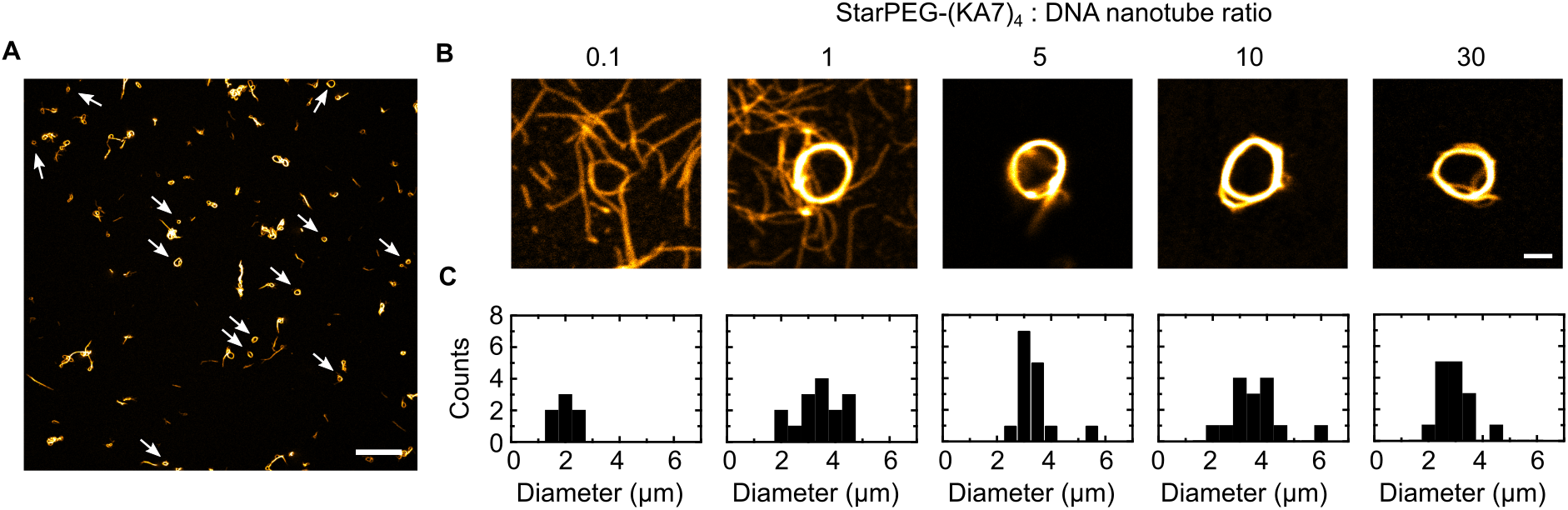
starPEG-(KA7)_4_ induces self-assembly of DNA nanotubes into micron-scale DNA rings. (**A**) Confocal overview image of self-assembled DNA nanotube rings formed from 50 nM DNA tiles (yellow, Atto633-labeled, *λ*_*ex*_ = 640 nm) in presence of 500 nM starPEG- (KA7)_4_. Rings are highlighted with white arrows. Scale bar: 20 µm. (**B**) Representative confocal images of individual DNA rings formed at starPEG-(KA7)_4_ to DNA nanotube (yellow, Atto633-labeled, *λ*_*ex*_ = 640 nm) ratios from 0.1 to 30. The DNA tile concentration is constant at 50 nM. Scale bar: 2 µm. (**C**) Histogram of the DNA ring diameters for starPEG-(KA7)_4_ to DNA nanotube ratios from 0.1 to 30 (*n* = 7 − 15).

Taken together, we obtain the first free-standing micron-scale DNA rings by self-assembly with synthetic crosslinkers. This mechanism mimics naturally occurring processes that have been shown to lead to the formation of actin rings (*24*).

### Contraction of DNA rings

Next, we set out to control the ring diameter and to induce ring constriction. It has been shown that rings formed from actin filaments can contract without consuming chemical energy (*4*). In this specific case, a crosslinker like annilin can induce entropy-driven filament sliding in order to maximize the overlap between filaments, which results in ring contraction (*4*). We observe, however, that the diameter of the DNA rings in presence of starPEG-(KA7)_4_ remains constant over time (Supplementary Fig. S9). This could be (i) due to the less tight linkage of the DNA nanotubes compared to actin and/or (ii) due to their greater persistence length. Hence we reason that addition of a molecular crowder could cause tighter linkage (i) and thus bring us into a regime where DNA ring contraction could occur. To test this hypothesis, we first form the DNA rings as previously in the presence of starPEG-(KA7)_4_. To induce tighter packing with molecular crowders, we add dextran to the DNA ring-containing solution. We find that the additional molecular crowding induces a DNA ring shrinkage by over 200 % from a mean diameter of (3.3 ± 0.7) µm to (1.4 ± 0.4) µm (Fig. 4**A**,**B**). Overall, the higher the molecular weight of dextran i.e. the higher the molecular crowding, the smaller is the ring size. Importantly, the rings’ shape, quantified by their circularity, remains unaltered and close to 1 (circularity of 1 is a perfect circle, Supplementary Fig. S10). We observe similar results with Methylcellulose as a molecular crowder (Supplementary Fig. S11). Importantly, the ring contraction shows that the crosslinking by starPEG-(KA7)_4_ and the additional molecular crowding keep the DNA nanotubes compliantly linked together. This allows dynamic structural changes that maximize the overlap between the DNA nanotubes, while inevitably reducing the ring diameter.

**Fig. 4.**
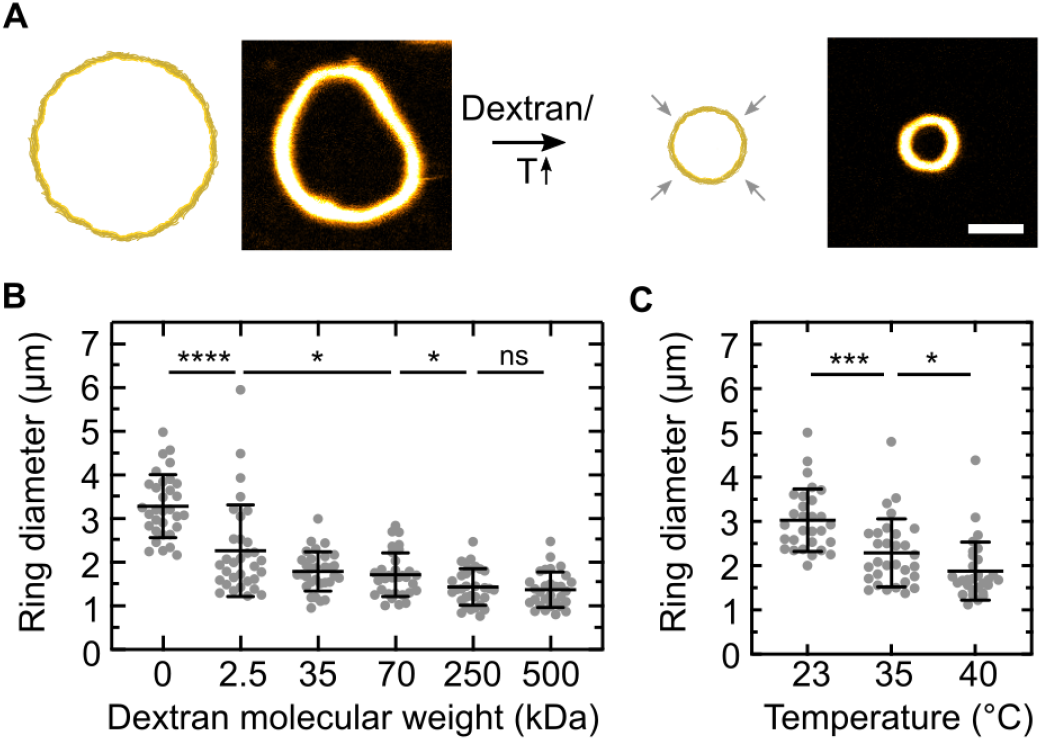
DNA nanotubes contract upon addition of a molecular crowder or heating. (**A**) Schematic representation and confocal images of DNA nanotube ring contraction (yellow, Atto633-labeled, *λ*_*ex*_ = 640 nm) in presence of 500 nM starPEG-(KA7)_4_. Dextran as a molecular crowder or temperature increase serve as triggers for contraction. Scale bar: 2 µm. (**B**) DNA nanotube ring diameter after addition of 25 wt% dextran with different molecular weights (Mean ± SD, *n* = 30 − 32). (**C**) DNA nanotube ring diameter after heating to 35 and 40 ° C, respectively (Mean ± SD, *n* = 29 − 31).

As an alternative to the addition of molecular crowders, we also achieve ring shrinkage due to temperature increase (Fig. 4**c**). This is especially desirable since this allows to control the ring contraction with an external trigger. Increasing the temperature from room temperature to 40° C leads to a reduction of the mean ring diameter from (3.0 ± 0.7) µm to (1.9 ± 0.7) µm. Note that the ring diameter does not change over time at constant temperature on the here relevant time scales (Supplementary Fig. S9). The temperature increase decreases the persistence length of the DNA nanotubes (ii) in such a way that the minimal energy conformation is shifted to smaller ring diameters. DNA rings thus contract significantly more than rings formed from actin filaments when they are contracted by passive crosslinkers (*4*). This is likely due to the lower persistance length of the DNA nanotubes compared to actin and thus their lower compliance. Both approaches for contraction, triggered by molecular crowding or temperature increase, yield a high degree of control over the ring size and allow the fine-tuning of ring diameters with more than a twofold change in ring diameter.

## Discussion

The reconstitution and stimuli-responsive control of the protein-based cellular division machinery has proven to be a difficult challenge. While it remains an attractive area of research, there is a quest for synthetic counterparts. Here, we engineer synthetic micron-scale DNA rings, self-assembled from a bundle of tens of DNA nanotubes and crosslinked via electrostatic interactions with custom-designed starPEG-(KA7)_4_ peptides. These micron-sized DNA rings could be used as tracks for molecular assembly and transport or embedded into adaptive materials. In addition, compared to their biological counterpart, synthetic DNA rings can easily be equipped with other types of molecular functionalization, reprogrammed and repurposed for diverse systems. Thus, micron-sized rings expand the toolbox for DNA assembly, in particular with DNA nanotubes. The possibility to contract the DNA rings with two different stimuli, molecular crowding and temperature increase, enhances their applicability for synthetic cellular systems. Nevertheless, an entirely synthetic division machinery for liposomes, based on DNA nanotechnology and peptide design, is a highly attractive albeit far-reaching goal. It has to be acknowledged that several challenges have to be overcome to induce vesicle contraction and division. The rings have to be positioned in the equatorial plane of the liposome, which will likely be achievable by self-assembly if the persistence length of the DNA nanotubes is sufficiently high (*12*). Secondly, the rings have to be linked to the membrane which could be achievable e.g. with cholesterol-tags (*12*). The temperature-induced contraction would in principle be compatible with liposome-encapsulation and the contraction force could potentially be sufficiently high to induce vesicle deformation (*23*). It could be complemented by engineered molecular motors that walk on DNA nanotubes (*25*). For ring disassembly, which will be necessary to complete the division of the compartment, it is plausible to use mechanisms that have already been described for DNA nanotubes (*11, 12, 15*).

We believe that the symbiosis of DNA nanotechnology and peptide engineering may lead to advanced and highly functional molecular hardware for bottom-up synthetic biology and hybrid materials with a wide range of applicability.

## Materials and Methods

### DNA Nanotube Design and Assembly

DNA nanotube sequences were adapted from the original single-tile design by Rothemund et al. (*26*). Each tile is composed of five DNA oligomers, the DNA sequences are listed in Supporting Table S1. The five DNA oligomers were mixed to a final concentration of 5 µM in 1× phosphate-buffered saline (PBS, pH 7.4) and 10 mM MgCl_2_. The tiles were annealed using a thermocycler (Bio-Rad) by heating the solution to 90 °C and cooling it to 25°C in steps of 0.5°C for 4.5 h. The assembled DNA nanotubes were stored at 4°C and used within two weeks. The DNA oligomers were purchased from Integrated DNA Technologies or Biomers (purification: standard desalting for unmodified oligomers, HPLC for oligomers with biotin and fluorophore modifications) and stored at 100 µM in 1× Tris-EDTA (pH 8) and stored at −20 °C.

### Peptide Synthesis

The synthetic peptides, KA7, DA7 and starPEG-(KA7)_4_, were synthesized as described in detail in (*23*). Briefly, all peptides were prepared by a standard Fmoc solid phase synthesis approach on an automated solid-phase peptide synthesizer (ResPep SL, Intavis) using 2- (1H-benzotriazole-1-yl)-1,1,3,3-tetramethyluronium hexafluorophosphate (HBTU) activation. Each amino acid was coupled twice with fivefold excess, while all non-reacted amino groups were capped with acetic anhydride. Likewise, 5(6)-TAMRA was coupled to the N-terminus of the peptides still bound the resin. Peptides were removed from the resin with trifluoroacetic acid/triisopropyl silane/water/dithiothreitol (DTT) (90[vol/vol]:5[vol/vol]:2.5[vol/vol]:2.5[m/vol]) for 1.5 h and precipitated with ice-cold diethyl ether. Peptides were further purified by reverse-phase high-pressure liquid chromatography (HPLC, Waters, Milford, MA) on a preparative C18 column (AXIA 100A, bead size 10 µm, 250 × 30 mm, Phenomenex, Torrance, CA) and analyzed by electrospray ionization mass spectrometry (ACQUITY TQ Detector; Waters) for purity. Cysteine-terminated KA7 and DA7 peptides were coupled to maleimide-terminated starPEG (10 kDa) by Michael addition reactions. For this, peptides and starPEG were mixed at a 1:5 (star-PEG:peptide) molar ratio in phosphate buffered saline (PBS, pH 7.4), sealed and stirred over night at 750 rpm and room temperature. The resulting star-PEG peptides were dialysed for 2 days against water using tubing with an 8-kDa cut-off. Peptide products were lyophilized and stored at −20 °C. For experiments, the respective peptides were rehydrated in 1× phosphate-buffered saline (PBS, pH 7.4) and stored as single-use aliquots at −20 °C. Peptide aliquots were stored at 1 mM and −20 °C. For experiments they were diluted in 1× phosphate-buffered saline buffer and used within one day.

### Confocal Fluorescence Microscopy

A confocal laser scanning microscope LSM 900 (Carl Zeiss AG) was used for confocal microscopy. The pinhole aperture was set to one Airy Unit and the experiments were performed at room temperature (unless stated otherwise). The images were acquired using a 20× (Plan-Apochromat 20×/0.8 Air M27, Carl Zeiss AG) or 63× objective (Plan-Apochromat 63×/1.4 Oil DIC M27). Images were analyzed and processed with ImageJ (NIH, brightness and contrast adjusted, ImageJ 2.3.0/1.5q; Java 1.8.0 322 64-bit, (*27*)).

### Charge, Crowding and Multivalency Control Assay

In order to compare the effect of different synthetic peptides on the DNA nanotubes, 30 nM DNA nanotubes were incubated with 500 nM positively charged multivalent starPEG-(KA7)_4_-TAMRA, 500 nM of the negatively charged multivalent starPEG-(DA7)_4_-TAMRA and 2 µM of the the positively charged monovalent (KA7)_4_-TAMRA or observed in absence of synthetic peptides. Buffer solutions contained 1× phosphate-buffered saline (pH 7.4) and 10 mM MgCl_2_. Samples were imaged in a custom-made observation chamber after one hour of incubation at room temperature (if not stated otherwise). The laser settings were kept the same at all time points. We analyzed the mean intensity per pixel in the same manner as we did for the colocalization assay. For each condition, nine overview images were analyzed, single data points and Mean±SD were plotted with Prism (GraphPad Prism Version 9.2.0).

### Colocalization Assay

For the preparation of DNA nanotubes A (green), the composition of DNA oligomer mix was altered to 90 % 6-FAM-labeled SE3 strand and 10 % biotinylated SE3 strand (Supporting Table 1). DNA nanotubes A were annealed as described above. 10 mg neutravidin was dissolved in 1 mL ultrapure water and then further diluted in PBS to a final concentration of 0.2 wt%. Biotinylated-BSA (Thermo Fisher) was diluted in ultrapure water to a concentration of 0.2 wt% and diluted in PBS to a final concentration of 0.1 wt%. The bottom glass slide of a custom-built observation chamber was coated with 0.1 wt% biotinylated BSA for 15 min. First, 50 µL of 0.2 wt% neutravidin were added on top of the slide and incubated for 2 min. After that, it was washed with PBS. Subsequently, 30 nM 6-FAM-labeled biotinylated DNA nanotubes A in 1× PBS and 10 mM MgCl_2_ were added. To seal the observation chamber, the upper slide was placed on top, spaced by double sided tape and sealed with two component glue. After 20 min incubation, the chamber was reopened and 50 µL of 5 µM starPEG-(KA7)_4_ diluted in 1× PBS was flushed. After an incubation of 2 min, 50 µL of 50 nM DNA nanotube B (green, Atto633-labeled) and diluted in PBS was flushed and washed again with 1× PBS. The observation chamber was sealed with two component glue for imaging. Images were acquired with confocal microscopy as described above.

For the starPEG-(KA7)_4_ colocalization assay increasing amounts of starPEG-(KA7)_4_-TAMRA were added to the DNA nanotubes. The mix contained starPEG-(KA7)_4_-TAMRA at varying concentrations (0, 50, 100, 200, 500 nM), 30 nM DNA nanotubes, 1× PBS and 10 mM MgCl_2_. The respective mix was left to incubate at room temperature for one hour. We then imaged the samples at the confocal laser scanning microscope with the 63x oil objective, keeping the same laser settings for all conditions.

We then analyzed the mean pixel colocalization intensity *c*_*p*_ of TAMRA-labeled peptides at the position of the nanotubes as follows. The analysis was performed with ImageJ (ImageJ 2.3.0/1.5q; Java 1.8.0 322 64-bit, (*27*)) and the plugin Skeleton. The pixel intensities for both images of either the DNA nanotube channel or the TAMRA channel range from *i* = 0 to 255 (Supplementary Fig. S2**A**,**D**). In order to extract the information about the DNA nanotube bundle positions, we first adjusted the threshold using the Method ‘Otsu’, thereby creating a binary image (Supplementary Fig. S2**B**). In a next step we used the plugin skeleton to reduce the DNA nanotube bundles to one single row of pixels in the middle of the DNA nanotube bundle, so we could generate a mask to always analyze the same area per nanotube length unit (Supplementary Fig. S2**C**). Then we linked the binary skeletonized information on location with the picture in the TAMRA-labeled channel by using the ImageCalculator function ‘AND’. Thereby we obtain background pixels counting zero intensity (*i* = 0) and skeletonized DNA nanotube (-bundle) pixels with non zero intensity (*i* = 1 to 255) (Supplementary Fig. S2**E**). The latter are included in the calculation of the average pixel intensity, called colocalization intensity. The resulting histogram carries the pixel number *p*_*i*_ for each intensity value *i* The pre-processing of the images directly leads to the fact that pixels with the intensity value zero are not included in the calculation of the average intensity at the DNA nanotube(-bundle) centers. The following equation outlines the relation.

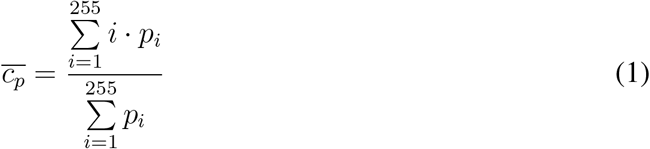

We calculated the mean pixel colocalization intensity *c*_*p*_ and standard deviation using ten overview images per sample. For 0 nM starPEG-(KA7)_4_ we analyzed five overview images. The plot was generated via Prism (GraphPad Prism Version 9.2.0 (332)). The same overview images were used to manually count the number of DNA rings.

### Transmission Electron Microscopy

For negative staining, 10 µL of DNA nanotube-containing solution (1× PBS, 10 mM MgCl_2_, 30 nM DNA nanotubes, and starPEG-(KA7)_4_ concentrations as described) 0.1 % paraformaldehyde was applied onto a glow-discharged 100 mesh copper grid with carbon-coated Formvar (Plano GmbH) and incubated for 30 min under a box. The solvent was removed by gentle blotting from one side with filter paper. The grid was rinsed with 3 drops of water, blotted again, and treated with 10 µL of 0.5 % (w/v) uranyl acetate solution for 20 s. After removing the staining solution thoroughly by blotting with filter paper, the grid was air dried and imaged on a FEI Tecnai G2 T20 twin transmission electron microscope (FEI NanoPort) operated at 200 kV. Electron micrographs were acquired with an FEI Eagle 4k HS, 200 kV CCD camera at 14500x nominal magnification.

### Bundle Thickness Analysis

30 nM DNA nanotubes were mixed with 0, 25, 50, 200 500 nM starPEG-(KA7)_4_ in 1× phosphatebuffered saline containing 10 mM MgCl_2_. Mixed samples were incubated at room temperature for 30 min and prepared for transmission electron microscopy. Images were acquired as described above at 14500x nominal magnification. Images were imported in ImageJ (ImageJ 2.3.0/1.5q; Java 1.8.0 322 64-bit, (*27*)) and the bundle thicknesses were evaluated by taking line profiles at manually selected positions for the DNA nanotube bundles. Per condition 100 positions were randomly chosen and measured. The data was plotted with GraphPad Prism 9 (Version 9.4.0) as Mean ± SD.

### StarPEG-(KA7)_4_-assisted DNA Ring Self-Assembly, Ring Size Control Assays and Ring Size Measurements

The starPEG-(KA7)_4_ stock solution (1 mM) was prediluted to 5 µM in PBS. The according amount of starPEG-(KA7)_4_ was added to the DNA nanotube mixture containing a final concentration of 50 nM DNA nanotubes in 10 mM MgCl_2_ and 1x PBS buffer (if not stated otherwise). The starPEG-(KA7)_4_:DNA nanotube ratio was set to 10 (if not stated otherwise). For colocalization experiments (Figs. 1**C**,**D**, 2**B**,**C**,**D**, Supplementary Figs. S2, S3, S5) and self-assembled ring observations (Fig. 3, Supplementary Figs. S6, S8, S9, S10, S11) 20 µL solution were added to a custom-built observation chamber. Self-assembled rings immediately stick to the uncoated glass slide due to the negative charge of DNA and are thereby immobilized for imaging.

For the ring size control with crowding molecules (Fig. 4**B**), 25 wt% Dextran (2 500, 35 000, 70 000, 250 000, 500 000 g*/*mol) was added to the DNA nanotube-starPEG-(KA7)_4_ mixture immediately after preparing the latter. Per condition, 100 µL of sample solution were prepared and entirely pipetted into a well slide (ibidi µ-Slide 18 Well, glass bottom) and imaged without delay at the confocal laser scanning microscope (Carl Zeiss AG) using the 63x oil objective. The laser settings were identical for all images.

For the ring size control with temperature increase (Fig. 4**C**), 500 µL of sample were prepared and at each time step 100 µL were pipetted into a well slide (ibidi µ-Slide 18 Well, glass bottom) inserted into a heating chamber (ibidi Temperature Controller, blue line). Lid and plate were set to the temperature of interest and the remaining sample was incubated at the same temperature for 15 min prior to pipetting.

For all ring size control experiments, 30 images of single rings were taken (5x zoom) per condition. Images were analyzed with ImageJ (ImageJ 2.3.0/1.5q; Java 1.8.0 322 64-bit, (*27*)) by tracing single rings with the freehand ROI selection. Circularity values were measured using the shape description feature of ImageJ. The perimeter *p* was measured, from which the diameter *d* was deduced by *d* = *p/π*. Thereby ring shapes that deviate from the circular were approximated to a circle and the ring size is expressed as the according diameter of an exact circle. Data was plotted as Mean±SD with GraphPad Prism 9 (Version 9.4.0).

### Statistical Analysis

All the experimental data were reported as Mean ± SD from *n* experiments, DNA nanotubes or rings. The respective value for *n* is stated in the corresponding figure captions. All experiments were repeated at least twice. To analyze the significance of the data, a Student’s t-test with Welch’s correction was performed using Prism GraphPad (Version 9.1.2) and p-values correspond to ****: p ≤ 0.0001, ***: p ≤ 0.001, **: p ≤ 0.01, *: p ≤ 0.05 and ns: p ≥ 0.05.

## Acknowledgements

K.G. acknowledges funding from the Deutsche Forschungsgemeinschaft (DFG, German Research Foundation) under Germany’s Excellence Strategy via the Excellence Cluster 3D Matter Made to Order (EXC-2082/1 – 390761711). M.I. and K.G. thank the Hector Fellow Academy. K.J. thanks the Carl Zeiss Foundation and the Joachim Herz Foundation for financial support. All authors acknowledge the Max Planck Society for its general support.

