## Supplementary Information for "Self-assembly and contraction of micron-scale DNA rings"

Illig, Jahnke *et al.*

and Kerstin Göpfrich,

#### Supplementary Figure S1: DNA bundles at high DNA nanotube concentrations

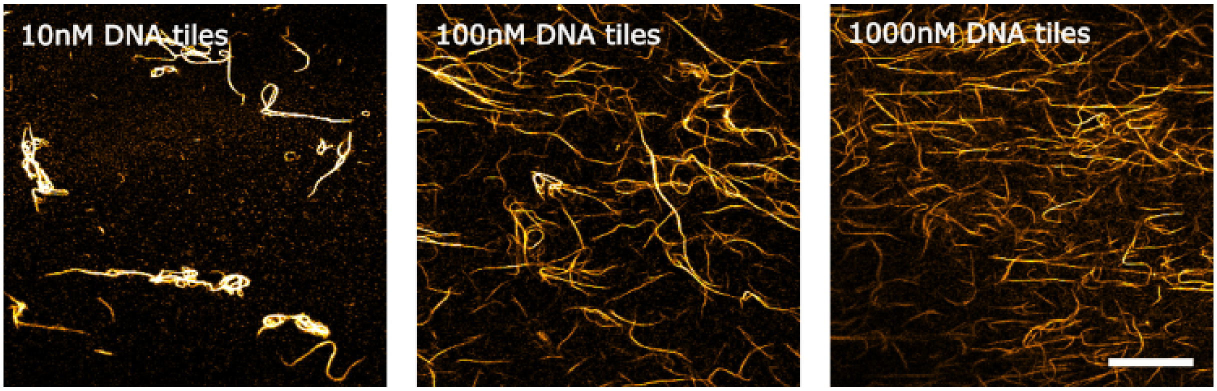


Fig. S1: DNA bundles at high DNA nanotube concentrations. Increasing amounts of DNA nanotubes (formed from $10\,\mathrm{nM}$, $100\,\mathrm{nM}$, $1000\,\mathrm{nM}$ DNA tiles) were incubated at room temperature with $500\,\mathrm{nM}$ starPEG-$(KA7)_4$-TAMRA in 1x PBS and $10\,\mathrm{mM}$ $MgCl_2$. The laser settings were kept the same at all conditions. Scale bar: $30\,\mu\mathrm{m}$.

#### Supplementary Figure S2: Colocalization intensity of TAMRA-labeled synthetic peptides

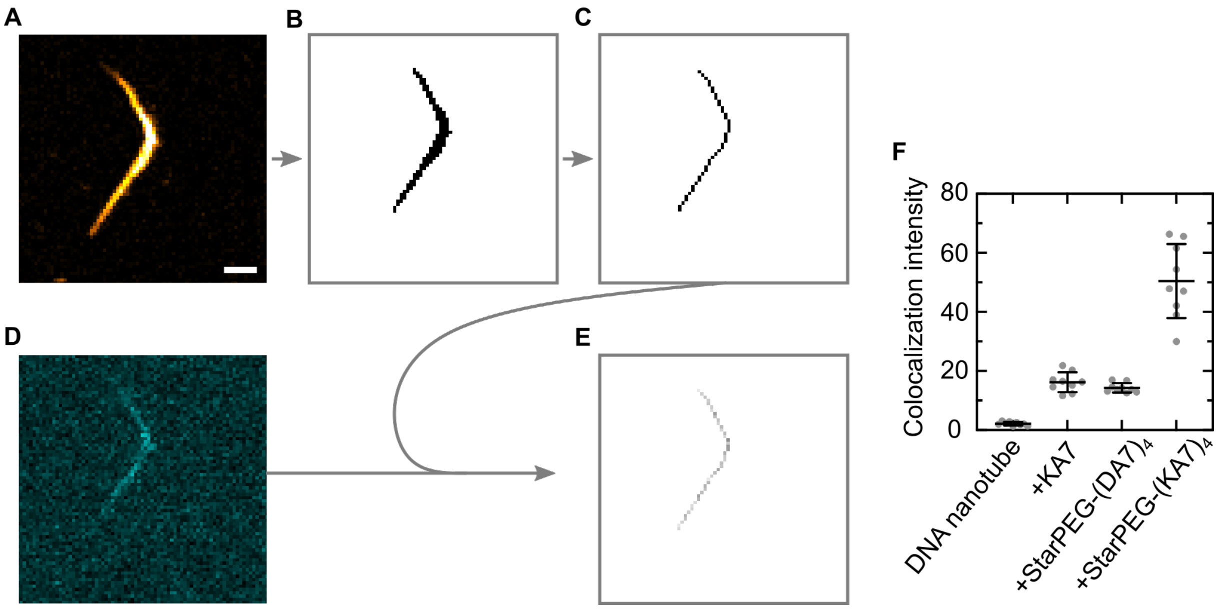


Fig. S2: Mean pixel colocalization intensity $c_p$ of TAMRA-labeled synthetic peptides. (**A**) Confocal image of $30\,\mathrm{nM}$ DNA nanotubes (yellow, Atto633, $\lambda_{ex} = 640\,\mathrm{nm}$) in presence of $500\,\mathrm{nM}$ starPEG-(KA7)$_4$ ($\lambda_{ex} = 561\,\mathrm{nm}$). Scale bar: $2\,\mu\mathrm{m}$. The image was analyzed with ImageJ ((Fiji Is Just) ImageJ 2.3.0/1.5q; Java 1.8.0_322 64-bit) (*1*) and thresholded via the method Otsu (**B**) and the binary image was processed further with the Plugin Skeletonize (**C**) to reduce the location of the nanotube bundle to the central line. Pixel values 0 and 1. (**D**) Confocal image of the same bundle in the starPEG-(KA7)$_4$ channel (cyan, $\lambda_{ex} = 561\,\mathrm{nm}$). Pixel values from 0 to 255. (**E**) starPEG-(KA7)$_4$ pixel intensity at the central line of the nanotube bundle, calculated by the ImageCalculator function 'AND'. Pixel values from 0 to 255. (**F**) Colocalization intensity of TAMRA-labeled synthetic peptides ($\lambda_{ex} = 561\,\mathrm{nm}$) with the DNA nanotubes (mean ± standard deviation (SD), $n = 9$). As expected, the colocalization intensity of starPEG-(KA7)$_4$ is the highest with 51.3 ± 12.5. The colocalization is significantly lower for KA7 and starPEG-(DA7)$_4$ (16.7 ± 3.3 and 14.1 ± 1.6, respectively).

#### Supplementary Figure S3: Colocalization of DNA nanotubes and starPEG

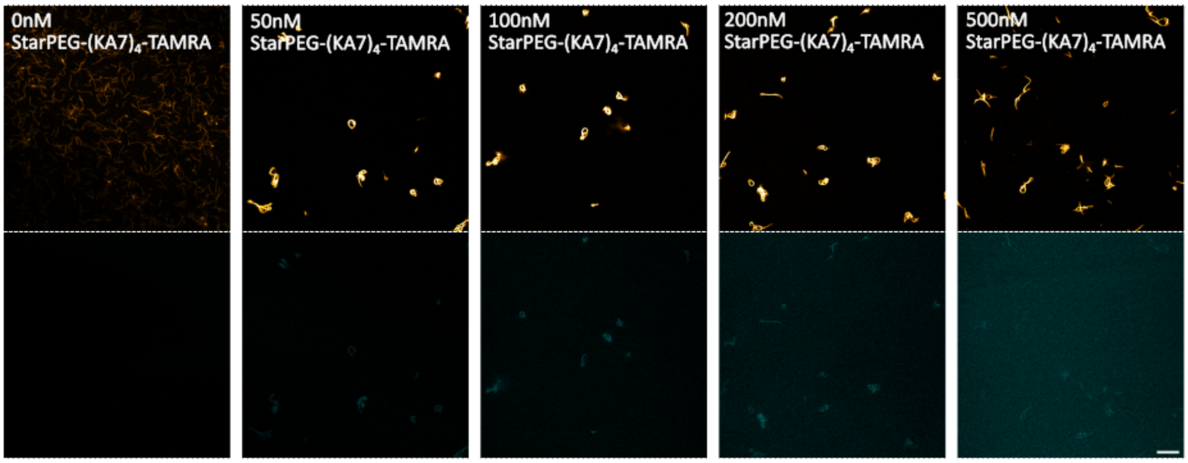


Fig. S3: Colocalization of DNA nanotubes and starPEG-$(KA7)_4$. The mix contained starPEG-(KA7)4-TAMRA at varying concentrations (0, 50, 100, 200, 500 nM), DNA nanotubes formed from 30 nM DNA tiles, 1× PBS and 10 mM $MgCl_2$. After incubation at room temperature for one hour the samples were imaged at the confocal laser scanning microscope with the same laser settings for all conditions. Scale bar: 10 μm.

#### Supplementary Figure S4: Transmission electron micrographs for bundle thickness analysis

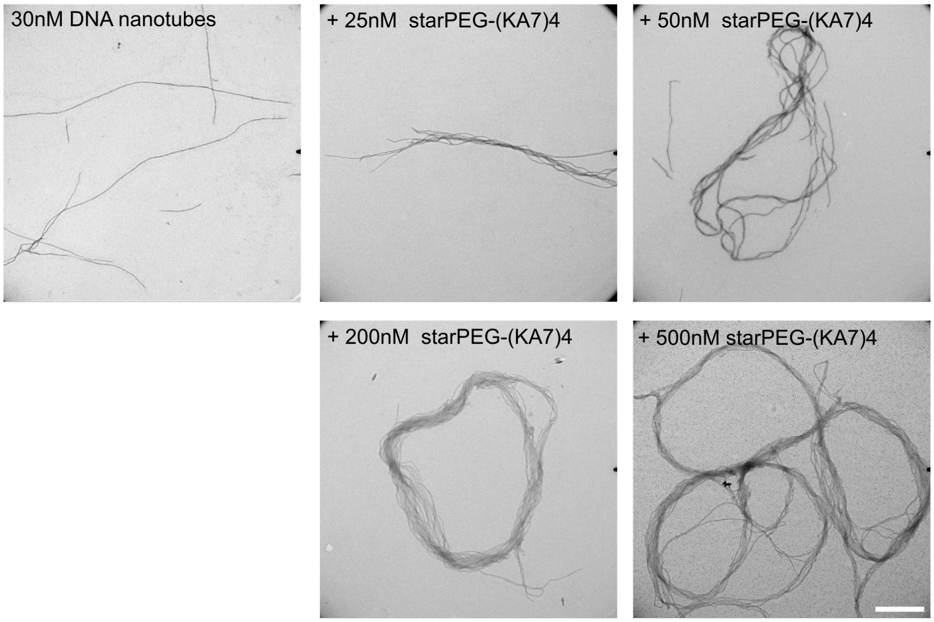


Fig. S4: Transmission electron micrographs of DNA nanotube bundles. $30\,\mathrm{nM}$ DNA tiles are bundled in the presence of $25$, $50$, $100$, $250$ and $500\,\mathrm{nM}$ starPEG-$(KA7)_4$ in 1× PBS and $10\,\mathrm{mM}$ $MgCl_2$. Bundles of different thicknesses were analyzed and the corresponding data is shown in Fig. 2**C**. Scale bar: $500\,\mathrm{nm}$.

#### Supplementary Figure S5: Colocalization intensity over time

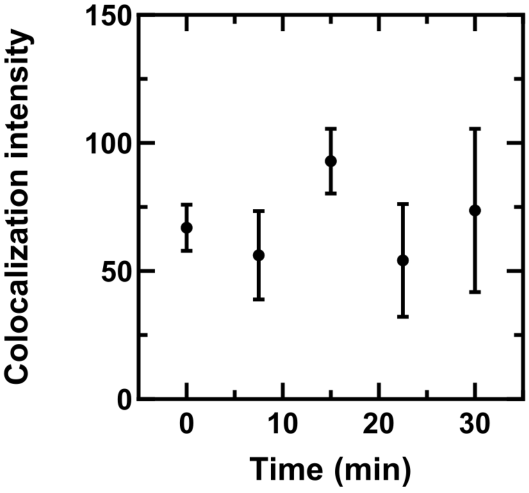


Fig. S5: Colocalization intensity over time. DNA nanotubes formed from $30\,\mathrm{nM}$ DNA tiles were left to incubate with $500\,\mathrm{nM}$ starPEG-(KA7)$_4$-TAMRA at room temperature. The sample was imaged at $0, 7.5, 15, 22.5, 30, 60$ and $120\,\mathrm{min}$ after adding $500\,\mathrm{nM}$ starPEG-(KA7)$_4$-TAMRA. The laser settings were kept the same at all time points. For each condition 10 overview images were analyzed (Mean$\pm$SD).

#### Supplementary Figure S6: Yield of DNA rings

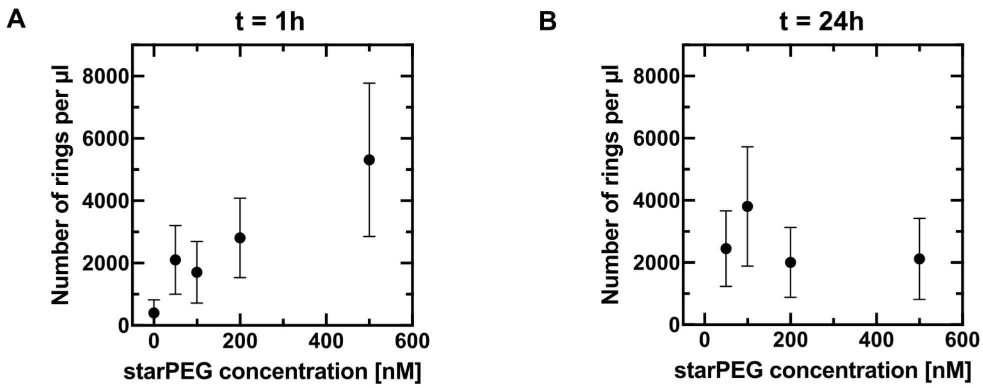


Fig. S6: Yield of DNA rings per µL for different starPEG-$(KA7)_4$ concentrations, $30\,\mathrm{nM}$ DNA nanotubes in 1× PBS and $10\,\mathrm{mM}$ $MgCl_2$. Per condition 10 overview images $(101.41\mathrm{x}101.41\,\mu\mathrm{m}^2)$ were acquired and rings were counted $1\,\mathrm{h}$ (**A**) and $24\,\mathrm{h}$ (**B**) after sample preparation. Volumetric densities were calculated by measuring the chamber height $(97\,\mu\mathrm{m})$ and taking all images into account (Mean$\pm$SD).

#### Supplementary Figure S7: Transmission electron micrographs of self-assembled DNA rings

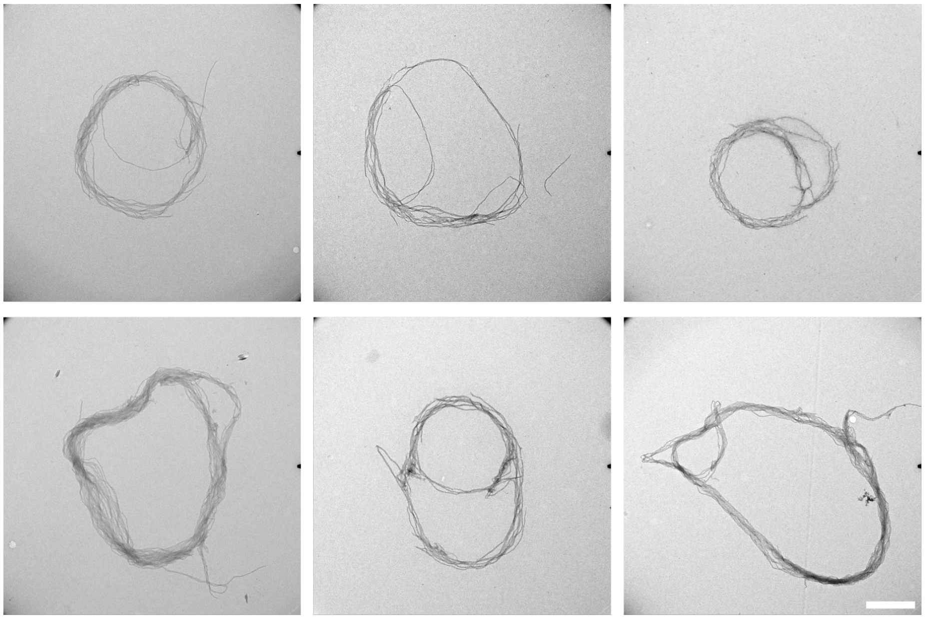

Fig. S7: Transmission electron micrographs of self-assembled DNA rings. $30\,\mathrm{nM}$ DNA tiles and $200\,\mathrm{nM}$ starPEG-$(KA7)_4$ are mixed in 1× PBS and $10\,\mathrm{mM}$ $MgCl_2$ and form rings of about 1-2 micrometer in diameter. Scale bar: $500\,\mathrm{nm}$.

#### Supplementary Figure S8: DNA ring size dependence on $MgCl_2$ and DNA nanotube concentration

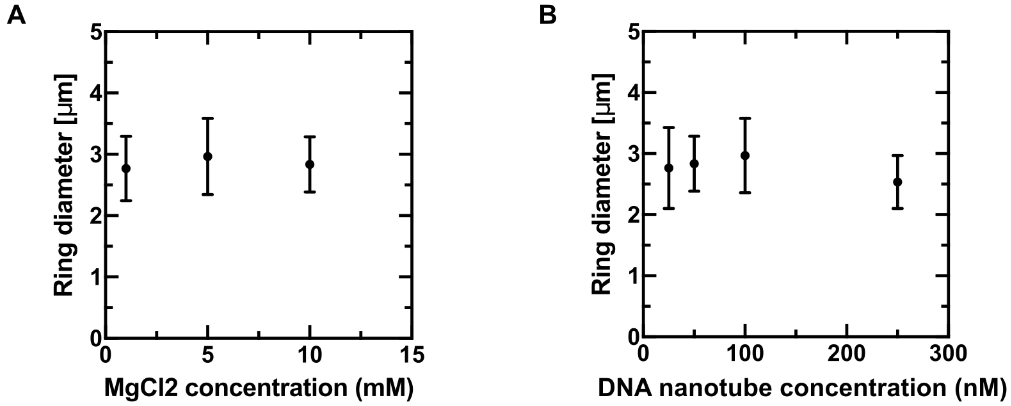


Fig. S8: DNA ring size dependence on $MgCl_2$ and DNA nanotube concentration. (**A**) Ring diameter at different buffer conditions containing $1$, $5$ or $10\,\mathrm{mM}$ $MgCl_2$ and $50\,\mathrm{nM}$ DNA tiles while maintaining the starPEG-$(KA7)_4$:DNA nanotube ratio of 10:1 ($500\,\mathrm{nM}$ starPEG-$(KA7)_4$) in 1× PBS. (**B**) DNA ring diameter for different DNA nanotube concentrations maintaining a starPEG-$(KA7)_4$:DNA nanotube ratio of 10:1 and $10\,\mathrm{mM}$ $MgCl_2$ and 1× PBS. Per condition 30 single rings were imaged and ring diameters are plotted as Mean$\pm$SD.

#### Supplementary Figure S9: DNA ring diameter is time-independent

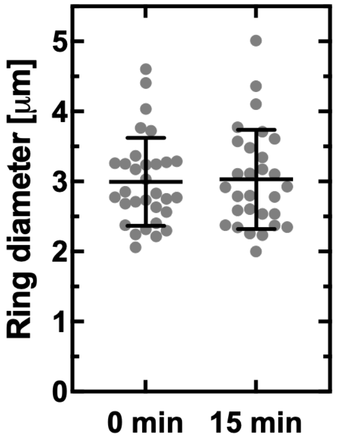


Fig. S9: DNA ring diameter is time-independent. DNA nanotube ring diameter at room temperature after $0$ and $15\,\mathrm{min}$ of incubation as used for the temperature increase experiments (Mean $\pm$ SD, $n = 29 - 31$).

#### Supplementary Figure S10: Analysis of the circularity of DNA rings

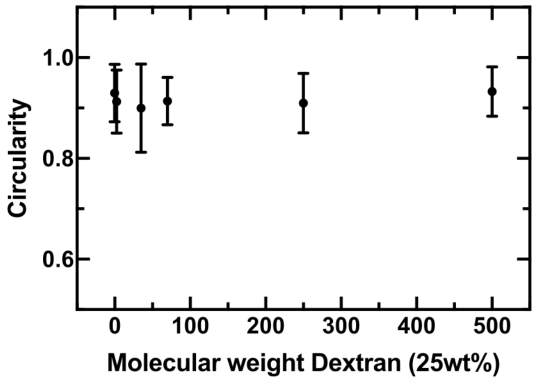


Fig. S10: DNA ring circularity for DNA nanotubes formed from $50\,\mathrm{nM}$ DNA tiles, $500\,\mathrm{nM}$ starPEG-$(KA7)_4$ in 1x PBS and $10\,\mathrm{mM}$ $MgCl_2$ and according molecular weights $(2\,500, 35\,000, 70\,000, 250\,000,\ 500\,000\,\mathrm{g/mol})$ of Dextran at $25\,\mathrm{wt\%}$.

### Supplementary Figure S11: DNA ring contraction with Methylcellulose

A

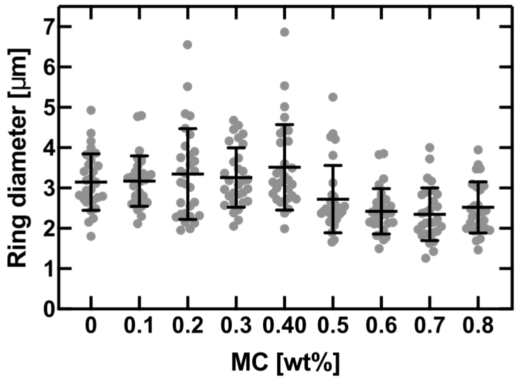


B

| | 0 | 0.1 | 0.2 | 0.3 | 0.4 | 0.5 | 0.6 | 0.7 | 0.8 |
|---|---|---|---|---|---|---|---|---|---|
| 0 | | Ns 0.8891 | Ns 0.8660 | Ns 0.6125 | Ns 0.2729 | ** 0.0028 | **** <0.0001 | **** <0.0001 | *** 0.0008 |
| 0.1 | | | Ns 0.8430 | Ns 0.7889 | Ns 0.3601 | *** 0.0007 | **** <0.0001 | **** <0.0001 | *** 0.0002 |
| 0.2 | | | | Ns 0.8660 | Ns 0.4232 | * 0.0389 | *** 0.0005 | *** 0.0001 | ** 0.0017 |
| 0.3 | | | | | Ns 0.4761 | *** 0.0007 | **** <0.0001 | **** <0.0001 | *** 0.0002 |
| 0.4 | | | | | | **** <0.0001 | **** <0.0001 | **** <0.0001 | **** <0.0001 |
| 0.5 | | | | | | | Ns 0.1529 | Ns 0.0561 | Ns 0.3637 |
| 0.6 | | | | | | | | Ns 0.5266 | Ns 0.6797 |
| 0.7 | | | | | | | | | Ns 0.2408 |
| 0.8 | | | | | | | | | |

Fig. S11: DNA ring contraction with Methylcellulose. $25\,\mathrm{nM}$ DNA nanotubes and $250\,\mathrm{nM}$ starPEG-(KA7)$_4$ are mixed in 1× PBS and $5\,\mathrm{mM}$ $MgCl_2$ and varied concentrations of Methylcellulose. Per condition, $100\,\mu\mathrm{L}$ sample solution is freshly mixed and pipetted into a well slide (ibidi μ-Slide 18 Well, glass bottom). (**A**) 30 rings are then imaged and analyzed using ImageJ (*1*) and the ring diameter is plotted (Mean±SD). (**B**) The ring diameter data from **A** is tested for significance. The corresponding p-values are obtained performing an unpaired non-parametric Mann-Whitney test.

#### Table S1: List of DNA sequences

| Name | DNA sequence |
|---|---|
| SE1 | CTCAGTGGACAGCCGTTCTGGAGCGTTGGACGAAACT |
| SE2-DIAG | GTCTGGTAGAGCACCACTGAGAGGTA |
| SE3 | CCAGAACGGCTGTGGCTAAACAGT<br>AACCGAAGCACCAACGCT(-6-FAM/Atto633/Biotin) |
| SE4-EE10 | CAGACAGTTTCGTGGTCATCGTACCT |
| SE5 | CGATGACCTGCTTCGGTTACTGTTTAGCCTGCTCTAC |

DNA sequences from 5’ to 3’ for single-tile DNA nanotubes, adapted from (*2*).
